# Limiting Self-Renewal of the Basal Compartment Induces Differentiation and Alters Evolution of Mammary Tumors

**DOI:** 10.1101/2020.01.18.907741

**Authors:** Nevena B. Ognjenovic, Meisam Bagheri, Gadisti Aisha Mohamed, Ke Xu, Meredith S. Brown, Youdinghuan Chen, Mohamed Ashick Mohamed Saleem, Shivashankar H. Nagaraj, Kristen E. Muller, Brock C. Christensen, Diwakar R. Pattabiraman

## Abstract

Differentiation therapy is an approach that utilizes our understanding of the hierarchy of cellular systems to pharmacologically induce a shift towards terminal commitment. While this approach has been a paradigm in treating certain hematological malignancies, efforts to translate this success to solid tumors have proven challenging. In this study we show that activation of PKA drives aberrant mammary differentiation by diminishing the self-renewing potential of the basal compartment. PKA activation results in tumors that are more benign, exhibiting reduced metastatic propensity, loss of tumor-initiating potential and increased sensitivity to chemotherapy. Analysis of tumor histopathology revealed features of overt differentiation with papillary characteristics. Longitudinal single cell profiling at the hyperplasia and tumor stages uncovered an altered path of tumor evolution whereby PKA curtails the emergence of aggressive subpopulations. PKA activation represents a promising approach as an adjuvant to chemotherapy for certain breast cancers, reviving the paradigm of differentiation therapy for solid tumors.

## Introduction

The process of differentiation involves a complex multi-step journey of cells from a state of multipotency towards one of commitment to a cell state that is, typically, post-mitotic. The process of tumorigenesis involves the acquisition of limitless proliferative capacity, the evasion of growth suppression, and the enabling of replicative immortality (Hanahan and Weinberg, 2011), all of which are characteristics of the circumvention of terminal differentiation. Differentiation therapy was first championed in hematological malignancies in which the use of all-trans-retinoic acid led to the terminal differentiation of acute promyelocytic leukemias (APL) (Huang et al., 1988; Warrell et al., 1993). Despite these advances, barring a few notable examples (Saha et al., 2014; Storm et al., 2016), differentiation therapy in solid tumors has largely been either unsuccessful or exploratory.

3’,5’-cyclic adenosine monophosphate (cAMP) is a widely used second messenger that regulates multiple downstream signaling cascades upon G-protein coupled receptor activation and subsequent Gα_s_ or Gα_i_-mediated regulation of adenylate cyclase (Tasken and Aandahl, 2004). One of the major routes of cAMP action is via activation of the ubiquitous enzyme Protein Kinase A (PKA). While in certain contexts, mutant G*α*_s_ acts in an oncogenic fashion (Patra et al., 2018; Wu et al., 2011), multiple lines of evidence suggest a role for Gα_s_-cAMP-PKA signaling in tumor-suppressive properties that could be mediated through the induction of differentiation (He et al., 2014; Iglesias-Bartolome et al., 2015; Pattabiraman et al., 2016).

Here we show that activation of PKA impairs mammary differentiation by specifically impairing self-renewal of the basal compartment. Mammary tumors with constitutively active PKA exhibit features of papillary differentiation with a reduced propensity for metastasis and increased susceptibility to chemotherapy. Longitudinal single cell profiling of mammary tumors at early and late stages of development revealed their inherent heterogeneity through the emergence of two subpopulations, lumino-basal, that co-express luminal and basal traits, and EMT-like, that have lost key epithelial markers. Both these subpopulations are severely diminished in the less aggressive PKA-activated tumors, implying that differentiation therapy strategies may favorably alter the representation of subpopulations that are responsible for tumor progression.

## Results

### Human Breast Cancers Harbor Recurrent Genomic Alterations in the PKA locus

Given the previously identified role for PKA in the induction of mesenchymal-epithelial transition (MET)(Pattabiraman et al., 2016), we interrogated the clinical significance of the PKA genes in human breast cancer. Analysis of the breast cancer genomic data from The Cancer Genome Atlas (TCGA)(Cancer Genome Atlas, 2012) revealed amplifications in the genomic loci encoding PKA subunits in 11.2% (121/1085) of breast cancers (Fig. 1A). Out of the 7 genes encoding PKA subunits, *PRKAR1A* and *PRKACA* were the two most frequently amplified across all tumors examined, while also being the two most highly expressed across all the molecular subtypes of breast cancer (Fig. 1B,C). *PRKAR1A* encodes one of four negative regulatory subunits that sequester the catalytic subunits, including that encoded by *PRKACA*, which carry out enzymatic functions downstream (Skalhegg and Tasken, 2000). Mining the METABRIC datasets (Curtis et al., 2012; Pereira et al., 2016) also revealed *PRKAR1A* amplifications in 11.9% (82/684) of tumors (Fig. 1D). In both datasets, amplifications were more frequent in the Luminal B subtype, with 13.03% and 14.32% of tumors in the TCGA and METABRIC datasets, respectively (Fig. 1D).

**Figure 1:**
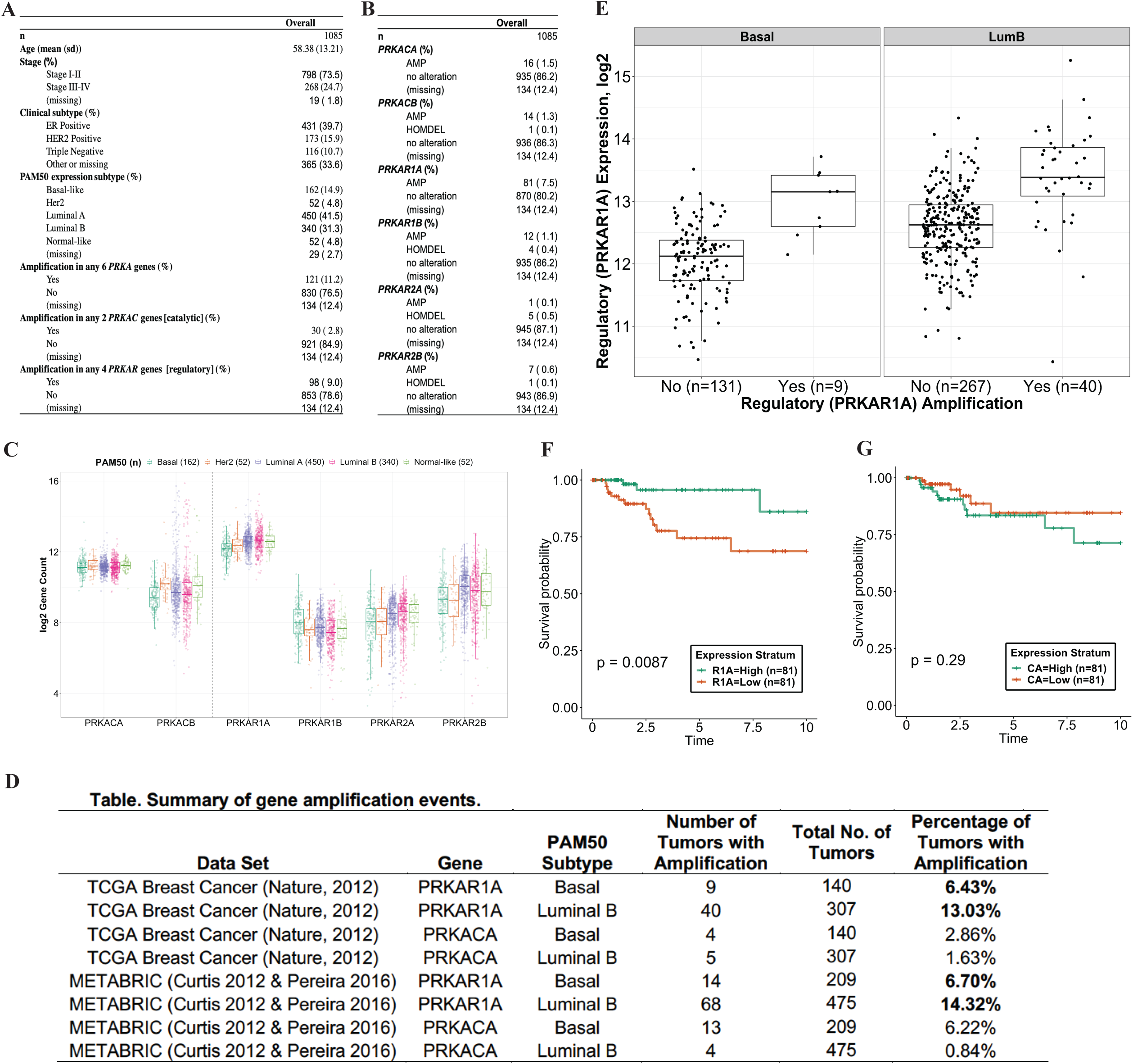
Recurrent genomic amplification of the *PKA* locus in human breast cancer. Analysis of 1085 breast tumors from the TCGA dataset revealing amplifications in the regulatory (*PRKAR*) and catalytic (*PRKAC*) genes that encode PKA subunits (A, B). (C) Box plots show the mRNA expression levels of genes encoding each PKA subunit and their relative expression levels across human breast cancer subtypes. Summary of *PRKAR1A* and *PRKACA* amplifications in the TCGA and METABRIC datasets in basal and luminal B subtypes (D). mRNA expression of *PRKAR1A*, the most amplified *PRKA* gene, across tumors with and without amplifications in the *PRKAR1A* locus (E). Kaplan-Meier curves outline the ability of high or low levels of *PRKAR1A* (F) or *PRKACA* (G) mRNA levels to stratify prognosis of patients over ten years.

*PRKAR1A* amplification correlated with higher expression levels of its mRNA (Fig. 1E), while also inversely correlating with expression levels of the catalytic subunit *PRKACA* (Fig. S1A,B) in a subtype-independent fashion. It was observed in the basal-like tumors of the TCGA dataset that higher *PRKAR1A* conferred better prognosis (Fig. 1F). These observations were, however, not reproduced in other datasets (Saal et al., 2015)(Fig. S1D,E), for other subtypes such as Luminal B (Fig. S1C), or for *PRKACA* (Fig 1G).

These recurrent genomic alterations in the PKA loci point to possible clinical implications for breast tumorigenesis, particularly of the luminal B subtype. We thus sought to study the role of activation of this pathway in mammary development and tumorigenesis.

### Constitutively Active PKA Signaling Impairs Mammary Differentiation

While previous studies have shown the ability of PKA to induce MET in some breast cancer cell lines (Pattabiraman et al., 2016), these data do not reveal the biological effects of PKA activation in mammary development and its impact on the subsequent development of tumors *in vivo*. We employed a mouse model harboring a floxed constitutively active mutant allele of the catalytic subunit of PKA (Niswender et al., 2005), and expressed this allele in the mouse mammary epithelium by crossing it to a mouse mammary tumor virus-driven Cre (*MMTV-Cre*) model (Wagner et al., 2001)(Fig. 2A). The mammary glands of these *Prkaca*^*CαR/fl*^ mice at eight weeks consistently contained fewer epithelial cells than littermate *Prkaca*^*fl/fl*^ controls, with a reduction of at least 50% in the EpCAM^med^CD49f^hi^ basal/myoepithelial cells (Fig. 2B,C), the compartment that harbors the highest proportion of mammary repopulating units. The postnatal mammary gland development in these mice was aberrant with a severe defect in ductal morphogenesis (Fig. 2D). In order to test whether the decrease in basal cells was merely quantitative, or whether activation of PKA led to a quantitative alteration of their self-renewal ability, we sorted for EpCAM^med^CD49f^hi^ basal cells from *Prkaca*^*CαR/fl*^ mice and control *Prkaca*^*fl/fl*^ littermates, and carried out organoid-forming assays (Guo et al., 2012). EpCAM^med^CD49f^hi^ cells from *Prkaca*^*CαR/fl*^ mice were severely impaired in the organoid-forming potential, indicating a loss of self-renewing potential (Fig. 2F,G). Additionally, when transplanted into the cleared fat pads of recipient mice, organoids from *Prkaca*^*CαR/fl*^ mice were at least 100-fold less efficient at forming ductal trees (Fig. 2E), revealing a loss of stem-like repopulating ability. This phenotype is reminiscent of previous observations in the skin whereby activation of Gα_s_-PKA signaling led to stem cell exhaustion and depletion (Iglesias-Bartolome et al., 2015).

**Figure 2:**
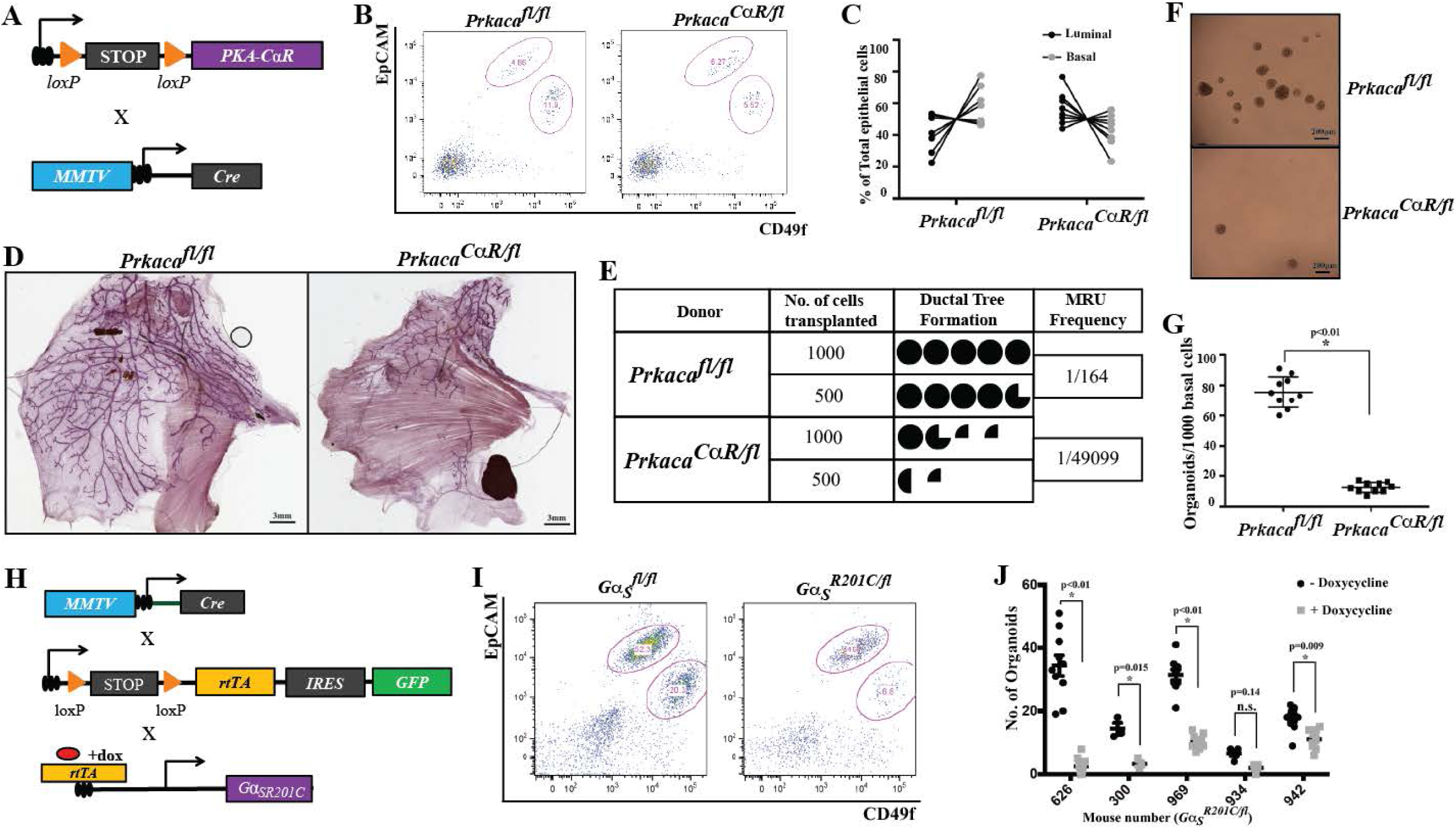
Activation of PKA impairs mammary development and repopulating ability. (A) PKA-CαR mice were crossed to MMTV-Cre mice to activate the *C*α*R* constitutively active allele. (B) FACS plots showing proportions of EpCAM^hi^CD49f^med^ luminal and EpCAM^med^CD49f^hi^ basal compartments upon constitutive activation of PKA and a summary of relative proportions of these subpopulations across multiple mice (C). Carmine alum stained whole mounts of mammary glands from and *Prkaca*^*CαR/fl*^ control mice displaying ductal outgrowth (D). Limiting dilution transplantation into cleared fat pads estimated the frequency of mammary repopulating units in *Prkaca*^*CαR/fl*^ and *Prkaca*^*fl/fl*^ controls (E). Organoid assays were carried out to assay for differences in self-renewal potential upon activation of PKA (F, G). Schematic representation of breeding strategy to generate mammary-specific *Gα*_*s*_ active mutants (H). FACS was carried out to capture differences in representation of luminal and basal subpopulations (I). Organoid assays carried out to test for differences in self-renewal potential (J). Error bars represent ± standard deviations of the mean. Statistical significance was calculated by a Student t-test (two-tailed) to compare two groups (P < 0.05 was considered significant).

In order to ensure that the effects observed could be attributed specifically to activation of PKA signaling, we used a second model, *Gα*_*s*_^*R201C/fl*^ mice (Iglesias-Bartolome et al., 2015), that express constitutively active *Gα*_*s*_ under the control of a doxycycline-inducible promoter. Upon expressing this allele in the mammary gland (Fig. 2H), doxycycline treatment was carried out to activate the *Gα*_*s*_*R201C* mutant from 3 weeks post-partum, at which point postnatal mammary development is initiated, to full maturity at 8 weeks. As observed in the *Prkaca*^*CαR/fl*^ mice, the *Gα*_*s*_^*R201C/fl*^ mice also exhibited a reduction in the EpCAM^med^CD49f^hi^ basal cells, which were severely impaired in their organoid-forming potential (Fig. 2I,J) while also displaying severe defects in mammary ductal outgrowth (Fig. S2A). Importantly, organoid-forming potential of basal cells from control *MMTV-Cre* mice was unaffected by doxycycline treatment (Fig. S2B).

To ensure that the observed results were not a result of MMTV promoter-mediated differences in the expression of the *CαR* allele in luminal vs. basal lineages, we crossed the *Prkaca*^*fl/fl*^ mice to a *R26-mTmG* reporter mouse (Muzumdar et al., 2007), and F1 mice were subsequently crossed with a *Krt14-CreERT2* mouse that would allow expression of the *C*α*R* allele in a basal-specific manner (Vasioukhin et al., 1999)(Fig. S2C). Upon tamoxifen administration at 4 weeks of age and observation of mammary glands at 8 weeks, we observed an absence of GFP-positive basal cells in the *Prkaca*^*CαR/fl*^ mice, present only in controls that lacked the *C*α*R* allele, or those that did not receive tamoxifen (Fig. S2D,E). These observations suggest that the expression of the *Prkaca*^*CαR*^ allele inhibited the expansion of the basal compartment, leading to the observed defects in ductal outgrowth. These results demonstrate the ability of PKA signaling in inducing a qualitative and quantitative alteration of the mammary basal/myoepithelial compartment.

We further explored the impact of these alterations on the development and progression of mammary tumors that arise from these aberrant glands.

### PKA-induced Differentiation leads to Attenuation of Metastatic Progression and Improved Response to Therapy

Given the high rate of recurrent *PRKAR1A* and *PRKACA* amplifications in the luminal B and basal subtypes of breast cancer (Fig. 1A), we chose to model PKA activation in the *MMTV-PyMT* mouse model of mammary tumorigenesis (Guy et al., 1992), which exhibit luminal B features in the early stages of tumor development, but acquire a more basal-like phenotype as tumors progress (Lin et al., 2003). Upon crossing the *Prkaca*^*CαR/fl*^ mice to the MMTV-PyMT mice (Fig. 3A), we observed that tumors in the control *Prkaca*^*fl/fl*^ PyMT mice took, on average, 142 days to reach the tumor volume endpoint of 2cm^2^, whereas those in the *Prkaca*^*CαR/fl*^ PyMT were slower to develop, taking 173 days (Fig. 3B), although overall tumor volume was comparable between the two strains (Fig. 3C). As expected, activation of the PKA signaling pathway was higher in the *Prkaca*^*CαR/fl*^ PyMT tumors compared to the *Prkaca*^*fl/fl*^ PyMT controls (Fig. 3D,E).

**Figure 3:**
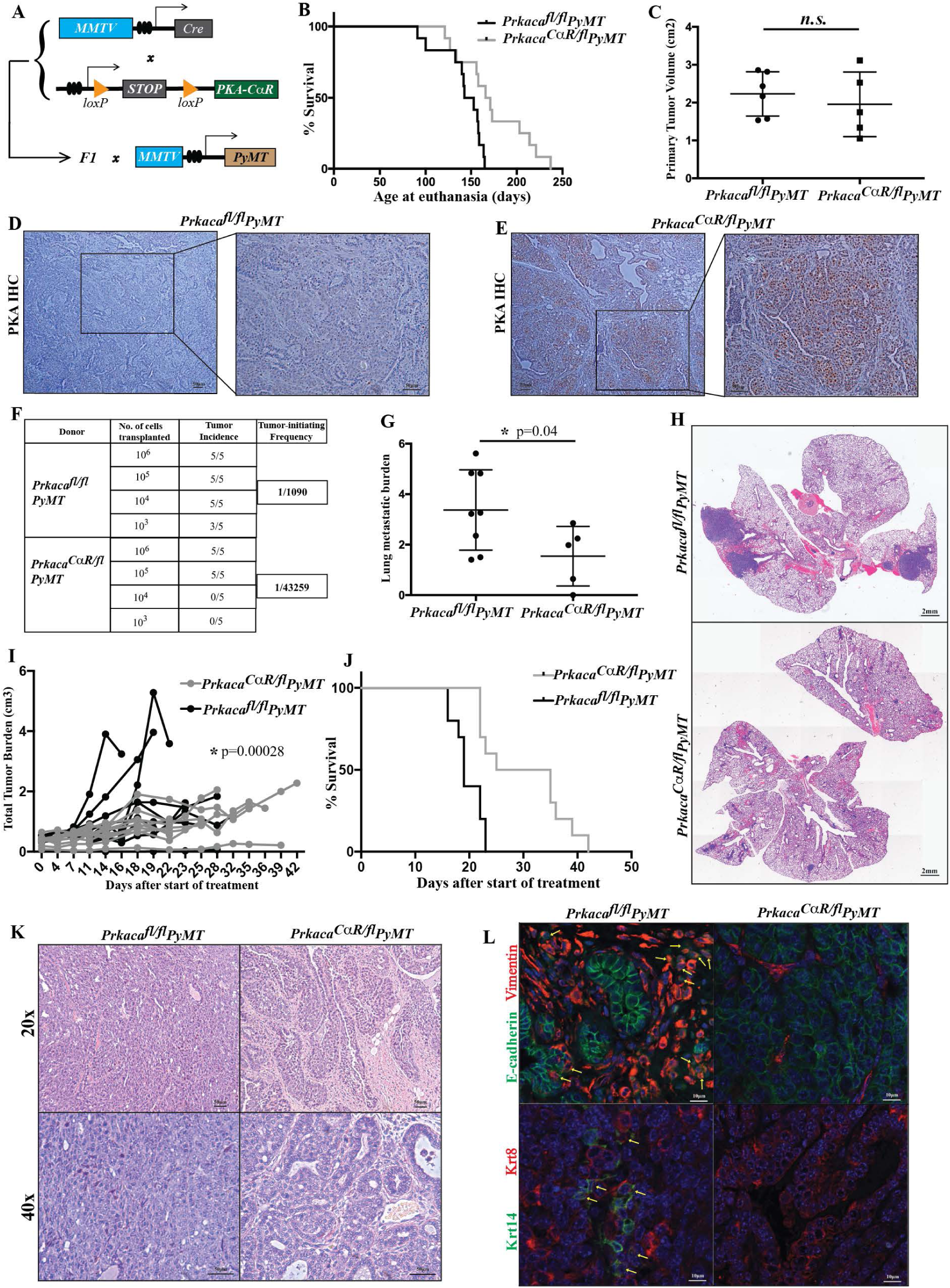
Induction of tumor differentiation upon activation of PKA signaling. (A) Schematic of breeding strategy to generate MMTV-PyMT mice with active PKA. Differences in survival (B) and primary tumor volumes (C) of *Prkaca*^*CαR/fl*^ PyMT mice compared to *Prkaca*^*fl/fl*^ PyMT controls. Activation of PKA was assessed by IHC using a p-PKA substrate antibody (D, E). Tumor initiating potential of *Prkaca*^*CαR/fl*^ PyMT mice and *Prkaca*^*fl/fl*^ PyMT controls was assessed by limiting dilution transplantation (F) analyses. Differences in lung metastatic burden was captured from H&E stained FFPE lung sections (G, H). Treatment with Adriamycin showed differences in response to chemotherapy (I, J) between *Prkaca*^*CαR/fl*^ PyMT mice and *Prkaca*^*fl/fl*^ PyMT controls. H&E staining of FFPE sections from the primary tumor was used to assess histopathology and differentiation state of the tumor (K), while IF using antibodies against E-cadherin, Vimentin, Krt8 and Krt14 revealed epithelial-mesenchymal and luminal-basal heterogeneity (L). Yellow arrows highlight double-positive cells. Error bars represent ± standard deviations of the mean. Statistical significance was calculated by a Student t-test (two-tailed) to compare two groups (P < 0.05 was considered significant). Significance in the chemotherapy treatment was established by a Wald Z-test was used to compute the p-value for the difference of slopes in two treatment groups.

Upon transplantation in limiting dilution into secondary hosts, *Prkaca*^*CαR/fl*^ tumors exhibited a 40-fold reduction in their tumor-initiating potential when compared to control tumors (Fig. 3F). In order to quantify the metastatic burden of the *Prkaca*^*CαR/fl*^ PyMT, we enumerated the number of lung macrometastatic foci from H&E stained sections and normalized this value to the primary tumor volume at the time of sacrifice. Thus, despite harboring primary tumors of comparable volumes (Fig. 3C) and having spent longer periods developing in the mice, the *Prkaca*^*CαR/fl*^ PyMT mice contained only half the metastatic burden of the *Prkaca*^*fl/fl*^ PyMT controls (Fig. 3G,H). Importantly, *Prkaca*^*CαR/fl*^ PyMT tumors were also more sensitive to treatment with adriamycin (Fig. 3I), a commonly used chemotherapeutic agent in the neoadjuvant treatment of human triple negative breast cancer. Treatment with Adriamycin significantly improved the survival of mice bearing *Prkaca*^*CαR/fl*^ PyMT when compared to controls (Fig. 3J).

Observation of hematoxylin-eosin (H&E) stained tumor sections revealed significant differences in their histopathology. The control *Prkaca*^*fl/fl*^ PyMT tumors resembled human invasive ductal carcinomas (IDC) of high grade comprised of solid sheets of pleomorphic cells with abundant mitotic activity (Fig. 3K). In contrast, tumors from *Prkaca*^*CαR/fl*^ PyMT mice exhibited a more differentiated phenotype with gland formation and blander nuclear characteristics, akin to an intermediate grade invasive ductal carcinoma. Additionally, these tumors contained a rich vascularized stroma reminiscent of papillary differentiation (Fig. 3K). Activation of PKA also resulted in a large reduction in tumor cells co-expressing Vimentin and E-cadherin (Fig. 3L) indicating a partial loss of EMT traits. The wild-type tumors, also harbored a significant number of Krt14-positive cells basal cells as previously reported (Cheung et al., 2013), a feature that was less prominent in *Prkaca*^*CαR/fl*^ PyMT tumors (Fig. 3L).

These histological differences in *Prkaca*^*CαR/fl*^ PyMT tumors could have occurred as a result of aberrant development of the mammary gland. To test whether induction of PKA in already formed tumors would have a similar effect, we used the inducible *Gα*_*s*_^*R201C/fl*^ model in which activation of PKA signaling would occur following treatment with doxycycline (Fig. S3A). Upon doxycycline induction at 8-9 weeks of age until sacrifice, *Gα*_*s*_^*R201C/fl*^ tumors also exhibited characteristics of papillary differentiation (Fig. S3B) indicating that the altered histological features were a result of activation of Gα_s_-PKA signaling. The induction of differentiation was also observed in a second model of mammary tumorigenesis, C3(1)-Tag (Maroulakou et al., 1994)(Fig. S3C), which develops basal-like tumors with features of metaplasia (Fig. S3D). *Prkaca*^*CαR/fl*^ C3(1)-Tag tumors also show signs of differentiation with a reduction in metaplastic and sarcomatoid features when compared to control *Prkaca*^*fl/fl*^ C3(1)-Tag tumors (Fig. S3D). Thus, the ability to induce differentiation is observed across different tumors types upon constitutive induction of PKA signaling. The resulting tumor pathology appears to depend on the nature of the tumor and its cellular origins.

### Differentiation Results from Altered Evolution of Tumor Subpopulations

In order to understand the molecular and cellular basis for the histopathological properties of the *Prkaca*^*fl/fl*^ PyMT and *Prkaca*^*CαR/fl*^ PyMT tumors, we longitudinally harvested cells at the early and late stages of tumorigenesis. At 9 weeks of age, the mammary glands of *Prkaca*^*fl/fl*^ PyMT mice are hyperplastic, while still largely maintaining their ductal architecture. We surgically resected one of two inguinal (no. 4) glands without euthanizing the animals, allowing them to develop tumors, which were later resected and sequenced. Using single-cell mRNA sequencing (scRNA-Seq) at both these time points, we were able to track the evolution of tumors at the hyperplasia and tumor stages (Fig. 4A). Data was analyzed using custom pipelines to pre-process, normalize and group the cells to identify cell (sub-) types that clustered together based on their gene expression patterns. Based on expression of well-known marker genes, clusters were annotated into basal, mature luminal, luminal progenitor, Lumino-basal and EMT-like (Fig. S4A-D).

**Figure 4:**
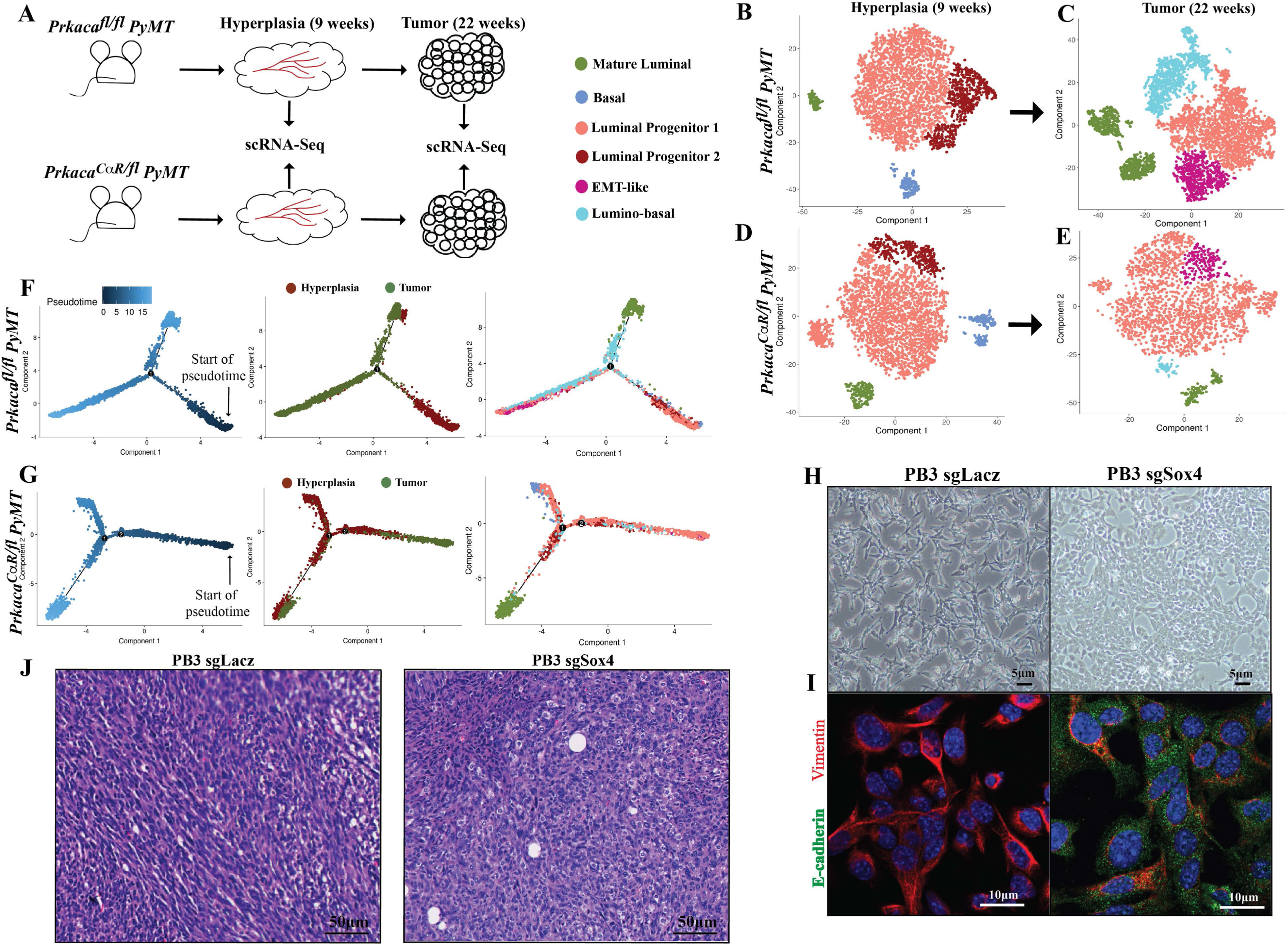
Altered evolution of tumor cell subpopulations upon activation of PKA. (A) Schematic of experimental set up to carry out sequential scRNA-Seq of hyperplastic and overt tumor samples from matched mice. tSNE plots show similar cellular constituents in hyperplastic glands of *Prkaca*^*CαR/fl*^ PyMT mice and *Prkaca*^*fl/fl*^ PyMT controls (B, D) but divergent evolution of subpopulations during the transition from hyperplasia to tumor stages of development (C, E). Pseudotime analyses enabled the capture of the directionality of evolution from hyperplasia to tumor in the *Prkaca*^*fl/fl*^ PyMT tumors (F) that is altered upon PKA activation in *Prkaca*^*CαR/fl*^ PyMT tumors (G). CRISPR-Cas9 mediated knockout of *Sox4* in PB3 cells leads to differences in cell morphology as captured by bright-field images (H) and the expression of epithelial-mesenchymal markers by immunofluorescence (I). H&E stained FFPE sections of transplanted PB3 tumors show differences in histopathology reflecting a change in differentiation state (J).

Dimensionality reduction and clustering of cells from *Prkaca*^*fl/fl*^ PyMT and *Prkaca*^*CαR/fl*^ PyMT mice yielded similar populations at the hyperplastic stage, with both exhibiting an expansion of the luminal progenitor compartment, while still retaining a portion of their normal basal and mature luminal subpopulations (Fig. 4B,D). There were, however, significant differences in the patterns of evolution from the hyperplasia to tumor stage. The *Prkaca*^*fl/fl*^ PyMT tumors harbored a luminal progenitor (LP) population that constituted the largest subpopulation within the tumor, but also contained a subpopulation, annotated lumino-basal (LB), which co-expressed LP and basal markers (Fig. 4C). These tumors also contained another subpopulation of cells, annotated EMT-like, that express relatively lower levels of a number of cytokeratins (Krt8, Krt14, Krt5, Krt79, Krt80), claudins (Cldn4, Cldn6, Cldn10), and differentiation markers (Prlr, Pgr, Lalba) (Fig. S4B,D). Notably, these subpopulations were evident only at the tumor stage, indicating that they were acquired between the two time-points sampled i.e., between the hyperplasia and tumor stages (9-22 weeks of age). Importantly, the *Prkaca*^*CαR/fl*^ PyMT tumors harbored significantly reduced populations of both the Lumino-basal and EMT-like clusters (Fig. 4G).

Given that cells from both stages of tumorigenesis were obtained from the same mouse, we carried out pseudotime analyses (Trapnell et al., 2014) to model the trajectories of tumor evolution. The kinetics of *Prkaca*^*fl/fl*^ PyMT tumors point to evolution from a hyperplastic signature to eventually acquiring a tumor signature (Fig. 4F). In contrast, the *Prkaca*^*CαR/fl*^ PyMT tumors exhibited an altered evolution trajectory whereby gene kinetics for tumor cells preceded that of hyperplastic cells in *Prkaca*^*CaR/fl*^ PyMT, with the tumor displaying a more differentiated profile than the hyperplasia (Fig. 4G). This provides a transcriptional basis for the *Prkaca*^*CαR/fl*^ PyMT tumors adopting a more differentiated histopathology, which likely drives the observed phenotypic and functional differences. The scRNA-Seq profiles reveal that the major cellular differences between the *Prkaca*^*CαR/fl*^ PyMT and controls lie in the evolution of the lumino-basal and EMT-like subpopulations. The presence of these cells drives the metastatic and therapy-resistant traits of the wild-type tumors, diminished in the PKA tumors. The emergence of these two subpopulations could mark the point at which wild-type tumors initiate metastatic spread.

### Sox4 is a Key Downstream Target of PKA-induced Differentiation

To further understand the emergence of the lumino-basal subpopulation, we looked to our scRNA-Seq data for the highest mRNAs expressed in the cluster. One of the key transcription factors that is highly expressed in the lumino-basal cluster is *Sox4* (Fig. S4E), a member of the Sox (SRY-related HMG-box) family of transcription factors that play important roles in embryonic development and cell-fate determination (Lioubinski et al., 2003). Other Sox family members have been shown to have profound impacts on mammary development and tumorigenesis (Dravis et al., 2015; Guo et al., 2012), whereas Sox4 has previously been demonstrated to play a role in EMT (Tiwari et al., 2013). In order to test whether Sox4 loss could recapitulate the effects of PKA activation, we carried out CRISPR-Cas9 mediated knockout of *Sox4* in PB3, a cell line derived from MMTV-PyMT tumors (Dongre et al., 2017). Loss of Sox4 induces a partial MET in PB3 cells, resulting in an altered morphology, upregulation of the epithelial marker E-cadherin and loss of Vimentin (Fig. 4H,I, Fig. S4F,G). Importantly, transplantation of Sox4^-/-^ PB3 cells results in the formation of tumors that exhibit characteristics of increased differentiation with more epithelioid cells in comparison with controls (Fig. 4J). Thus, loss of Sox4 phenocopies at least certain properties of the MET induction previously attributed to PKA (Pattabiraman et al., 2016). The partial nature of this phenotype suggests that the phenotypic effects of PKA induction are mediated through a number of downstream mechanisms, of which Sox4 is a part.

## Discussion

While differentiation therapy in solid tumors has had limited success, this study provides evidence for the utility of the PKA pathway as a differentiation-inducer in some breast tumors. This strategy may also be applicable to other tumor types where PKA has been shown to induce stem or progenitor cell differentiation, such as those driven by epidermal and hair follicle stem cells (Iglesias-Bartolome et al., 2015), in the adrenal cortex (Drelon et al., 2016), and granule neural progenitors (He et al., 2014).

While our previous studies revealed a role for PKA in altering EMT state of breast cancer cell lines, the effects of doing so on mammary development and the subsequent emergence of tumors, were unknown. For the first time, we show that activation of PKA leads to aberrant mammary development. The prevention of ductal outgrowth stems from a defect in basal/myoepithelial cell self-renewal, which also hinders the repopulation of ducts upon transplantation. Features of papillary differentiation are observed in tumors that arise from glands that had constitutive activation of PKA (Fig. 3K), as well as upon activation of PKA following initiation of tumorigenesis (Fig. S3B). Interestingly, the lack of an intact myoepithelial layer is a feature of malignant papillary carcinomas (Pal et al., 2010), indicating that the resulting papillary features could be a direct effect of a decrease in myoepithelial cells. Moreover, the activation of Gα_s_-PKA signaling appears to confer papillary features in other tumor-types (Wu et al., 2011), although the overall tumorigenic or tumor suppressive effects are tissue-type dependent.

Previous studies identified a subpopulation in MMTV-PyMT tumors that expresses basal markers and is responsible for collective invasion (Cheung et al., 2013). Our scRNA-Seq results also point to the emergence of a subpopulation of cells, the lumino-basal (LB) fraction, that is correlated with increased metastatic burden. We observe that these cells still retain the expression of luminal markers, including Krt8 and CD14, while also expressing basal markers. Our scRNA-Seq analyses reveal that activation of PKA suppresses the acquisition of basal traits that appear to be necessary to generate LB cells, mirroring the effects observed in the normal mammary gland.

The presence of recurrent genomic amplifications in *PRKAR1A* suggests a role for suppression of this pathway in tumorigenesis. Our results indicate that the catalytic subunit of the complex, encoded by *PRKACA*, plays a critical role in differentiation through its enzymatic activity. This suggests that higher levels of *PRKACA* could confer better prognosis in breast cancer patients, whereas the negative regulators of the catalytic subunit, encoded by *PRKAR1A* and other genes, could confer poorer prognosis. Our results, paradoxically, point to *PRKAR1A* conferring better prognosis in TCGA dataset in basal-like tumors (Fig. 1E). It must be noted that this does not translate to other datasets such as METABRIC, or even to other subtypes such as Luminal B, which harbor the majority of amplifications. The prognosis of tumors likely depends on the interactions of the protein products of these genes, rather than their mRNA levels that were used to stratify patients. Moreover, these data do not take into account the intratumoral heterogeneity present within tumors whereby minor subpopulations could dramatically influence outcomes. Nevertheless, it is conceivable that the negative regulation of this catalytic subunit by higher levels of the regulatory subunit could prime tumor cells to undergo epithelial-mesenchymal transition (EMT) and contribute to tumor progression.

Our study provides proof-of-principle for the utility of the induction of differentiation as a means to deriving therapy responsiveness in solid tumors, while curtailing their aggressive traits and metastatic spread. The ability of constitutively active PKA signaling to deplete self-renewal of basal cells in multiple tissue types suggests that careful calibration of its activity is essential for the maintenance of the stem cell state and the regulation of cellular differentiation. While targeted and immunotherapies have shown immense success in a number of cancer types, the inherent heterogeneity of some breast cancers are a major barrier to the identification of recurrent genomic alterations that are targetable. For such cancer types, a more feasible approach could be the reversal of a trait that is common to all cancers: the evasion of terminal differentiation. The induction of differentiation through activation of PKA, or through inhibition of key targets such as Sox4, could play an important adjuvant role to conventional chemotherapy to obtain better clinical outcomes.

## Materials and Methods

### Plasmids

pLentiCRISPRv2 was a gift from the Feng Zhang via Addgene (#52961, http://n2t.net/addgene:52961; RRID:Addgene_52961). sgRNAs for Sox4 were cloned into the construct using protocols from the Zhang lab. The Sox4 sgRNA sequences that were used were:

sgSox4_2: CAACAACGCGGAGAACACTG

sgSox4_4: CGACAAGATTCCGTTCATCC

sgLacZ: TGCGAATACGCCCACGCGAT

### Cells

PyMT tumor-derived PB3 cells were obtained from the Robert A. Weinberg lab and were cultured in DMEM/F12 medium containing 5% adult bovine serum with 1× penicillin– streptomycin and 1× nonessential amino acids. For the Sox4 knockout studies, cells were cultured for 14 days in the presence of puromycin for complete CRISPR-Cas9 mediated loss of Sox4, following which they were used for subsequent studies including immunoblotting, immunofluorescence and transplantation.

### Mammary Gland Preparation

Mouse mammary fat pads (no. 3 and 4) were harvested and dissociated as per previously published protocols(Prater et al., 2013) with minor modifications. Briefly, glands were digested with Collagenase A and Hyaluronidase for 2-3 hours at 37°C in a rotator followed by mild trituration with a 10ml pipette. Following red cell lysis, cells were treated with trypsin, and subsequently with dispase to aid dissociation. Cells were then stained with antibodies (see table below) to analyze and sort luminal and basal populations

### Carmine Alum Staining

Harvested mammary glands are spread out on a slide and allowed to dry for 1 hour at RT. Slides are fixed overnight in Carnoy’s fixative (6 parts 100% EtOH, 3 parts CHCL3, 1 part glacial acetic acid) at RT. On day 2 slides were rehydrated by incubating in 70% ethanol twice for 10 min each, twice in 50% ethanol for 10 min each, twice in 30% ethanol for 10 minutes each, twice in 10% ethanol for 10 minutes each and in distilled water for 5 min. Slides are then stained for 1-2 days in Carmine Alum (Stem Cell Technologies; Catalog No. 07070) at RT. Slides are dehydrated by incubating in 70% ethanol twice for 10 min each, twice in 95% ethanol for 10 min each and twice in 100% ethanol for 10 minutes each. Slides were then cleared in Xylene for 3-4 days, with the solution being changed every day after which they are rehydrated and dehydrated again as above before mounting using Permount (Fisher Chemical; SP15). Slides were scanned on a Perkin Elmer^©^ Vectra 3 slide scanner and analyzed on Phenochart^©^.

### Organoid Assays & Transplantation

Dissociated mammary epithelial cells were dissociated into single cells and cultured with advanced DMEM (Gibco; 12491015) containing 5% Matrigel, 5% heat-inactivated FBS, 10 ng/ml EGF, 20 ng/ml bFGF, 4 mg/ml heparin, and 5 mM Y-27632. Cells were seeded at 1,000 per well in 96-well ultralow attachment plates (Corning; 29443-034). The number of organoids was counted 7–14 days after seeding.

1000 or 500 cells from dissociated organoids were resuspended in 10 μl 75% DMEM/25% matrigel were injected into the number 4 glands of 4w old NSG female mice that had been cleared of endogenous epithelium. Recipient glands were harvested at 12wk, dissected, fixed and stained with carmine alum. An outgrowth was defined as an epithelial structure comprising ducts and branchings and the percentage of outgrowth was quantified by proportion of ducts and branches in relation to the fat pad.

### Animal studies

Research involving animals complied with protocols approved by the Dartmouth College Committee on Animal Care. The *PKA-CαR* (Niswender et al., 2005) mice were a gift from Stanley McKnight (UWashington). The *Gα*_*s*_R201C mice (Iglesias-Bartolome et al., 2015) were a gift from J. Silvio Gutkind. The B6.Cg-Gt(ROSA)26Sor^tm1(rtTA,EGFP)Nagy/J^ mice (Belteki et al., 2005) (Stock No. 005670), mTmG reporter mice - B6.129(Cg)-Gt(ROSA)26Sor^tm4(ACTB-tdTomato,-EGFP)Luo/J^ mice (Muzumdar et al., 2007) (Stock No. 007676), Tg(KRT14-cre/ERT)^20Efu/J^ mice (Vasioukhin et al., 1999) (Stock No. 005107), Tg(MMTV-PyVT)^634Mul/LellJ^ mice (Guy et al., 1992) on a C57Bl/6J background (Stock No. 022974) and Tg(C3-1-TAg)^cJeg/JegJ^ mice (Maroulakou et al., 1994) (Stock No. 013591) were obtained from The Jackson Laboratory. For transplantation studies, cells suspended in DMEM containing 30% Matrigel (GFR)/PBS mix (BD Biosciences; 356230) were injected into the inguinal mammary gland fat pads of age-matched female NOD/SCID or C57Bl6/J mice (Jackson Laboratory). Mice were euthanized after 10 weeks or when tumors reached a diameter of 2 cm. For chemotherapy studies, mice were administered adriamycin intraperitoneally at 2mg/kg twice a week for up to 40 days or until tumors reach the endpoint of 2cm in diameter.

### Tumor Dissociation

Tumors from MMTV-PyMT mice and C3(1)-Tag mice were resected and minced using a razor blade in DMEM containing 2 mg/mL collagenase and 100 U/mL hyaluronidase (Roche) in a rotator at 37°C for 30 minutes. Following incubation, tumors were minced further and re-incubated for another 30 minutes. Dissociated cells were washed twice in PBS and filtered through a 70- and 40-μm cell strainer to obtain single-cell suspensions.

### Immunofluorescence (cultured cells)

Cells were cultured on dishes containing coverslips for 2 to 3 days, after which coverslips were washed in cold PBS, fixed in 5% Neutral Buffered Formalin for 10 min at 4°C and permeabilized in 0.2% TritonX in PBS for 2 min. Cells were then washed in PBS, blocked for 1 hour at room temperature in PBS containing 3% normal horse serum (Vector Labs, USA; S-2000). Fixed cells were then incubated with the primary antibody in PBS containing 1% bovine serum albumen (BSA) solution overnight at 4°C. Cells were washed in PBS three times, and secondary antibody was added in PBS containing 1%BSA solution for 1 to 2 hours at room temperature in the dark. Cells were washed three times in PBS and were incubated for 2min in DAPI solution, after which they were washed in PBS and mounted with a drop of Prolong Diamond antifade reagent (Life Technologies; P36962) and placed on coverslips. Slides were viewed on a Zeiss^©^ LSM800 microscope and analyzed using the Zen^©^ Digital Imaging software.

### Immunofluorescence (FFPE tissue sections)

Slides were rehydrated by incubating in Histoclear solution twice for 5 min each, followed by incubation in 100% ethanol twice for 5 min each, in 95% ethanol twice for 5min each, 70% ethanol twice for 5 min each, once in 35% ethanol for 5 min, and in water for 5 min. Pressure cooker– mediated heat-induced epitope retrieval was carried out in 250 ml of unmasking buffer containing sodium citrate at pH 6. After retrieval, slides were blocked for 30 min in PBS containing 3% normal horse serum after which they were incubated with primary antibody in blocking solution overnight at 4°C. Slides were washed twice with PBS and incubated with secondary antibody at room temperature for 1 hour in the dark. After two PBS washes, 20 ml of mounting medium was added, then slide contents were topped with coverslips, and stored in the dark for 24 hours before imaging on a Zeiss LSM800 microscope and analyzed using the Zen Digital Imaging software.

| **Antibody** | **Vendor** | **Catalog No.** | **Application** | **Dilution** |
| --- | --- | --- | --- | --- |
| CD45 PECy7 | BioLegend | 103114 | FACS | 1:100 |
| CD31 PECy7 | BioLegend | 102418 | FACS | 1:100 |
| EpCAM APC | BioLegend | 118214 | FACS | 1:100 |
| CD49f PE | BioLegend | 313612 | FACS | 1:100 |
| Vimentin | Cell Signaling | 5741 | IF/IHC | 1:100 |
| E-cadherin | BD | 610182 | IF/IHC, WB | 1:200 |
| Keratin 8 | Thermo Fisher | PA5-29607 | IF/IHC | 1:400 |
| Keratin 14 | Thermo Fisher | MA5-11599 | IF/IHC | 1:1000 |
| Sox4 | Thermo Fisher | MA5-31424 | WB | 1:1000 |
| p-PKA substrate | Cell Signaling | 9624 | IHC | 1:200 |

### Analysis of publicly available datasets

The Cancer Genome Atlas (TCGA) (Cancer Genome Atlas, 2012) and METABRIC (Curtis et al., 2012; Pereira et al., 2016) breast cancer gene expression, copy number, clinical and survival data, were accessed from the cBioPortal database (www.cbioportal.org). The Swedish Cancerome gene expression and clinical data were publicly available in the Gene Expression Omnibus database (www.ncbi.nlm.nih.gov/geo/) under accession GSE96058 (Saal et al., 2015). All statistical analyses were performed in the R statistical computing environment (v3.5.1). Specifically, the correlation between mRNA expression of two genes were assessed by ordinary least-square regression. The univariate Kaplan-Meier analyses and related data visualizations were implemented in R packages *survival* (v.2.43.3) and *survminer* (v.0.4.3)(Therneau and Grambsch, 2000).

### Single-cell RNA-seq library construction and sequencing

Single Cell Capture and Library Preparation: Immediately following dissociation, single cell suspensions were placed on ice and counted on a Luna automated cell counter. Cell concentrations from each sample were be normalized to 1000 cells/ul and loaded onto a Chromium Single Cell A Chip (10x Genomics Inc.) targeting a capture rate of 5,000 cells per sample. Single cell RNA-seq libraries were be prepared using the Chromium Single Cell 3’ v2 kit (10x Genomics) following the manufacturer’s protocol. Libraries were quantified by qubit and peak size determined on a fragment analyzer instrument. All libraries were pooled and sequenced on an Illumina NextSeq500 High Output 26bp × 98bp run to generate an average of 50,000 reads/cell. Data Analysis: Raw sequencing were processed using the 10x Genomics Cell Ranger to generate quality metrics and primary data visualizations as well as gene expression matrices for downstream analysis in R using Seurat and other open-source packages.

### Single-cell RNA-seq data processing

To investigate the transcriptional evolution of tumors at single cell level, we performed of matched hyperplasia and tumor samples was performed on the 10X genomics platform, and generated data at an average of ∼78M reads per sample with cell numbers ranging from ∼2600 to 5000. Single-cell transcriptome sequencing raw reads were quality filtered using *Fqtrim* (v0.9.7) tool [10.5281/zenodo.1185412]. Reads were trimmed and filtered for low quality bases, poly-A/T tails and N bases while retaining the paired-end integrity of the reads. Read1 of the pair containing barcodes was not considered for trimming but allowed to be filtered to maintain paired-end integrity. Sequencing and PCR errors in cell barcodes can convolute the process of differentiating reads per cell barcode; hence we used *UMItools* (v0.5.4) to distinguish the reads per cell (barcode) accommodating for technical errors(Smith et al., 2017). A knee density method-based approach is used in *UMItools* to estimate the number of acceptable cell barcodes and then reads were assigned for each cell.

Clean and barcode classified reads were aligned against Mouse genome reference (GRCm38) using *STAR* aligner (v2.5.3a)(Dobin et al., 2013) and output was restricted to uniquely aligned reads. De-duplication of transcripts with the same UMIs arising from PCR amplification were removed. Reads were assigned with position based annotation of genomic features using *featureCounts* module(Liao et al., 2014) in the *subread* package (v1.6.0). Taking advantage of the UMI information, read counts were extrapolated to quantify molecular level count for each transcript using directional-adjacency method-based count module in *UMItools*. Reads were grouped per cell (based on barcode) and then a gene expression matrix was generated with RNA molecule count of genes in rows for each cell in columns represented in GxC matrix format (where G is gene in rows and C is cells in columns).

### Cell-type classification and Pseudotime analysis

Gene expression matrix was filtered for cells with <500 genes expressed and <500 total UMI counts genes and >0.25 percentage of reads aligned to Mitochondrial genome using *Monocle2* (v2.6.4) R package(Qiu et al., 2017a; Qiu et al., 2017b; Trapnell et al., 2014). Further, outliers of total mRNAs count for each cell was removed from the downstream analysis. Expression counts were normalized using negative binomial distribution of library sizes. Genes with mean expression value of 0.1 across cells were used for PCA based dimensionality reduction and then clustered using an unsupervised *densityPeaks* algorithm in *Monocle2* and projected using *t-SNE* method.

To classify the population into various cell-types, we investigated the expression of curated and well-established cell-type specific marker genes’ expression in our data and selected a specific list of markers that are expressed and/or not expressed for each cell-type. Following are the cell-type specific markers used for the classification in this study,

| Cell types | Higher expression |
| --- | --- |
| Basal cells | *Krt14, Krt5, Snai2* |
| Luminal Mature cells | *Pgr, Prlr, Foxa1, Gpx3, Esr1* |
| Luminal Progenitor cells | *CD14, Kit, Lalba* |
| Lumino-basal | *Krt14, Krt8, CD14* |
| EMT-like | *Trim29, Vim, Cldn6, Sox4* |

We identified and classified cell-types using *Monocle2*’s *classifyCells* function. First, each cell was annotated into a cell-type based on the presence and absence of positive and negative cell-type marker genes respectively. Then cells were re-clustered using added cell-type information to bring cells with similar gene-expression pattern and cell-type annotation into closer clusters, and to segregate un-annotated cells. Ambiguous and Un-annotated clusters were investigated manually for other known marker genes using *find_cluster_markers* function in *Monocle3 alpha* (v2.99.3) and re-annotated manually into different sub-types based on GO enrichment analysis of selected marker genes and expression of other known cell-type specific marker genes.

Pseudotime analysis attempts to reconstruct the transcriptional transitions of cells during the process of tumorigenesis. By predicting the transcriptional state of each cell, cells are ordered into an estimated pseudotime indicating their cell-state and transition across a time-series. Transition of gene expression kinetics was performed by integrative Pseudotime analysis using *Monocle2*. Given that the hyperplasia and tumor stage cells were extracted from the same mice, the hyperplasia and tumor stage samples of same mice were merged using *Seurat* R package (v2.3.0)(Butler et al., 2018) while retaining the cell-type and sample annotation. Merged data was processed again for dimensionality reduction using *DDRTree* algorithm and then pseudotime trajectories were calculated. Plots were generated using different color codes based on cell-type, sample and pseudotime to illustrate the gene expression kinetics transition across stages and samples. Data plots were generated using *Monocle2* and *Monocle3 alpha* packages.

## Statistical analysis

Data are presented as means ± SD. A Student’s t test (two-tailed) was used to compare two groups (P < 0.05 was considered significant) unless otherwise indicated.

## Author Contributions

**Conception and design**: D.R.P

**Development of methodology**: N.B.O, M.B, G.A.M, K.X, Y.C, M.A.M.S, S.H.N, B.C.C, D.R.P

**Acquisition of data**: N.B.O, M.B, G.A.M, K.X, M.S.B, D.R.P

**Analysis and interpretation of data**: N.B.O, M.B, G.A.M, K.X, Y.C, M.A.M.S, S.H.N, K.E.M, B.C.C, D.R.P

**Writing, review, and/or revision of the manuscript**: D.R.P

**Study supervision**: D.R.P

## Acknowledgements

We thank Dr. Stanley McKnight for the *PKA-C*α*R* mice and Dr. Silvio Gutkind for the *Gα*_*s*_*R201C* mice; the genome technology core at the Whitehead Institute and the Genomics and Molecular Biology Shared Resource at the Norris Cotton Center for optimization of protocols for single cell RNA sequencing; Dartlab, Microscopy, Pathology and Mouse Modeling Shared Resources at the Norris Cotton Cancer Center. We thank Ferenc Reinhardt, Joana Liu Donaher, Jennifer Fields and Rebecca O’Meara for technical assistance and Drs. Yashi Ahmed and Alan Eastman for critical reading of the manuscript. Funding and resources for the shared resources were supported in part by a core grant (5P30CA023108-40; Norris Cotton Cancer Center). This work was supported by funding from the NIH R01CA216265 (to B.C. Christensen) and 5R00CA201574-05 (to D.R. Pattabiraman).

## Supplementary Figure Legends

**Supplementary Figure 1:**
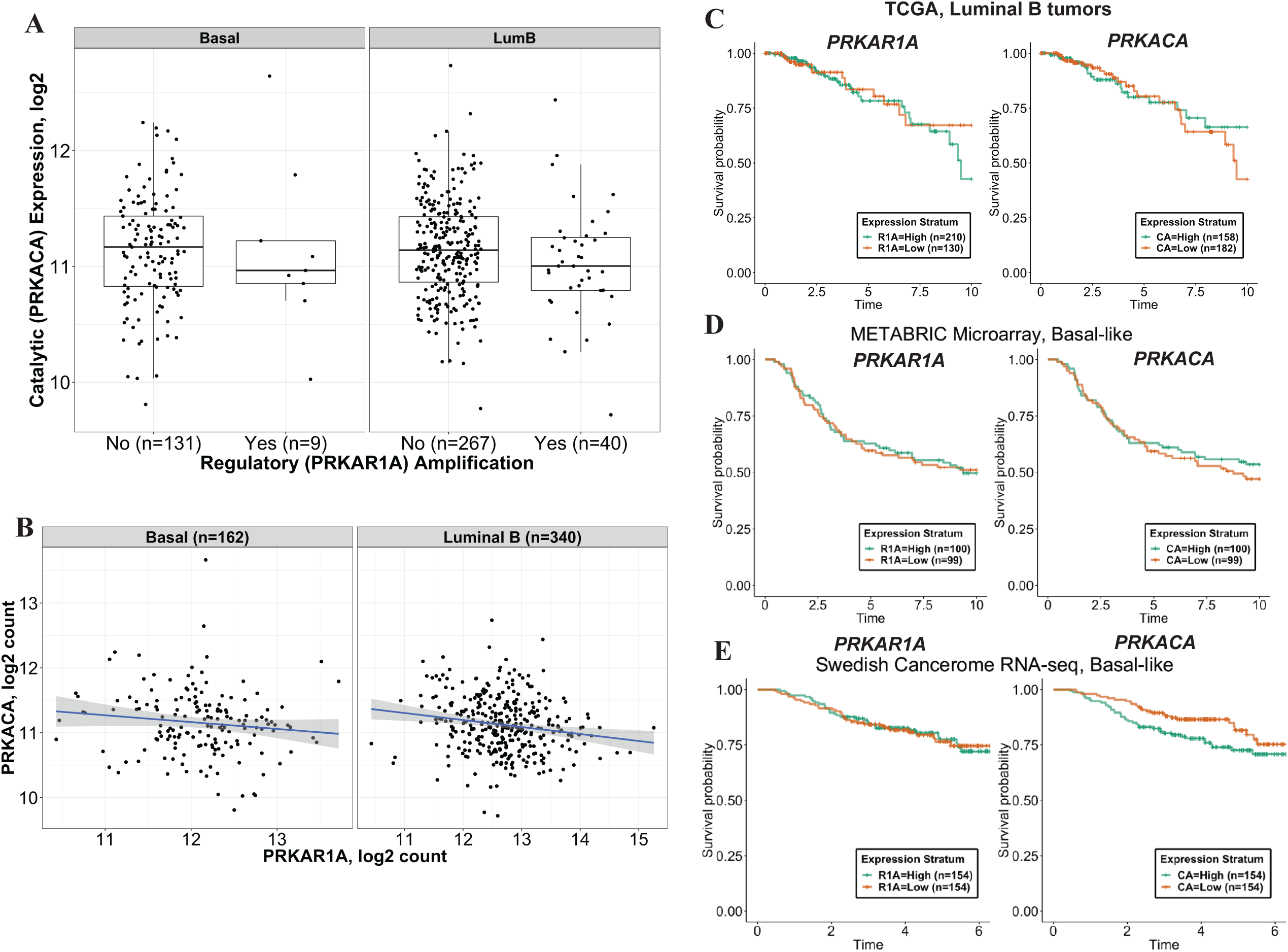
(A) Scatter box plots representing gene expression levels of *PRKACA* in basal and luminal B breast cancers that do or do not harbor amplifications in the *PRKAR1A* locus. Results are from the TCGA dataset (B) Scatter plots showing correlation between expression of *PRKACA* and *PRKAR1A* across basal and luminal B tumors of the TCGA dataset. Kaplain-Meier curves reveal the ability of expression levels of *PRKAR1A* and *PRKACA* to stratify patients based on prognosis in (C) luminal B tumors from the TCGA dataset, (D) Basal-like tumors from the METABRIC dataset and (E) Basal-like tumors from the Swedish Cancerome RNA-Seq dataset.

**Supplementary Figure 2:**
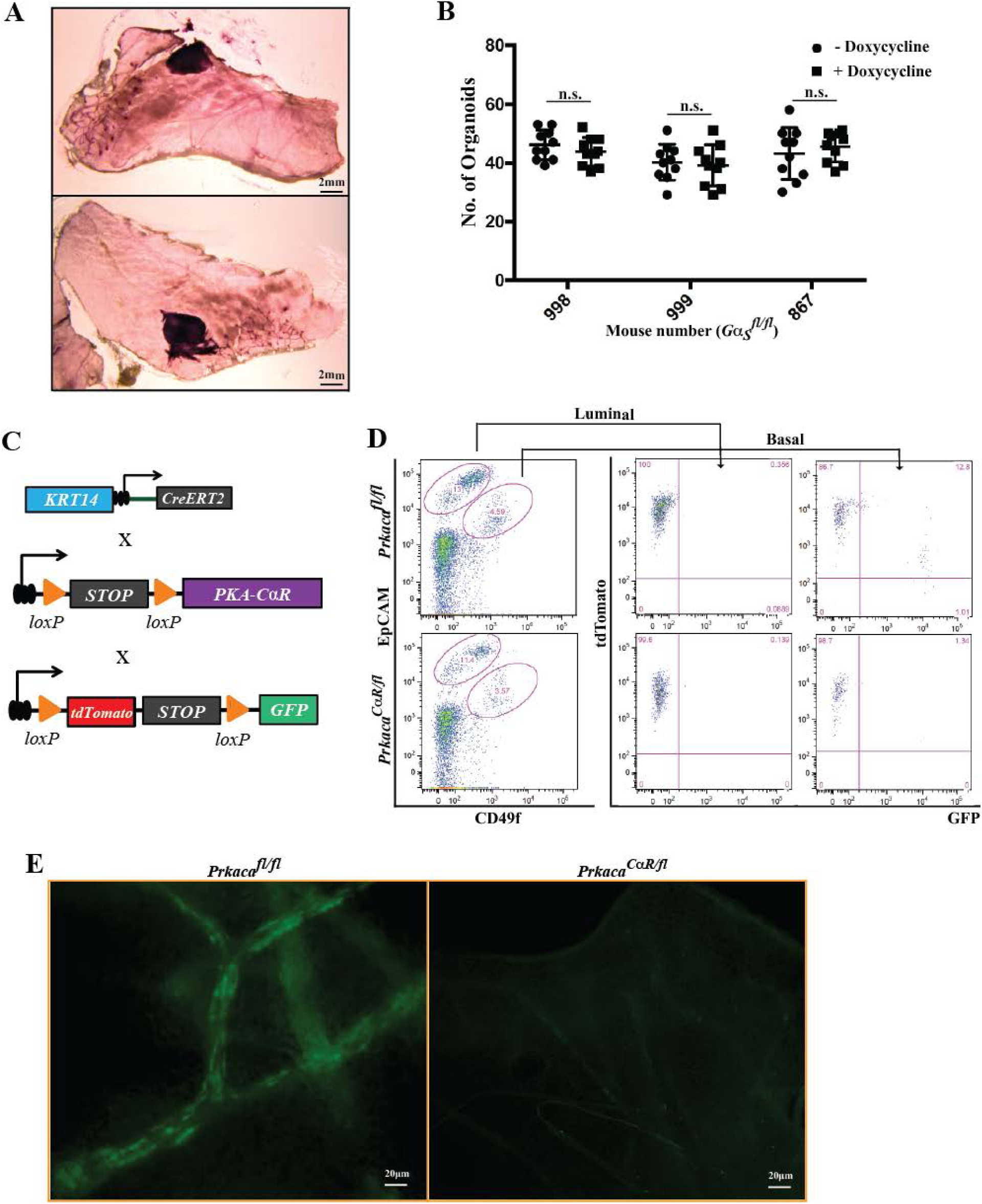
(A) Scatter plot outlines the number of organoids enumerated from the culture of EpCAM^med^CD49f^hi^ basal cells from *Gα*_*s*_^*fl/fl*^ mice from three individual mice in the absence and presence of doxycycline. (B) Whole mount images outlining the morphology of carmine alum stained mammary glands from *Gα*_*s*_^*R201C/fl*^ mice. (C) Schematic of mouse crossing strategy to label and express the *PKA-C*α*R* allele specifically in basal cells, following which FACS analysis (D) and fluorescence imaging (E) was carried out on glands that were harvested at 8 weeks to assess the effects of the allele in the mammary development.

**Supplementary Figure 3:**
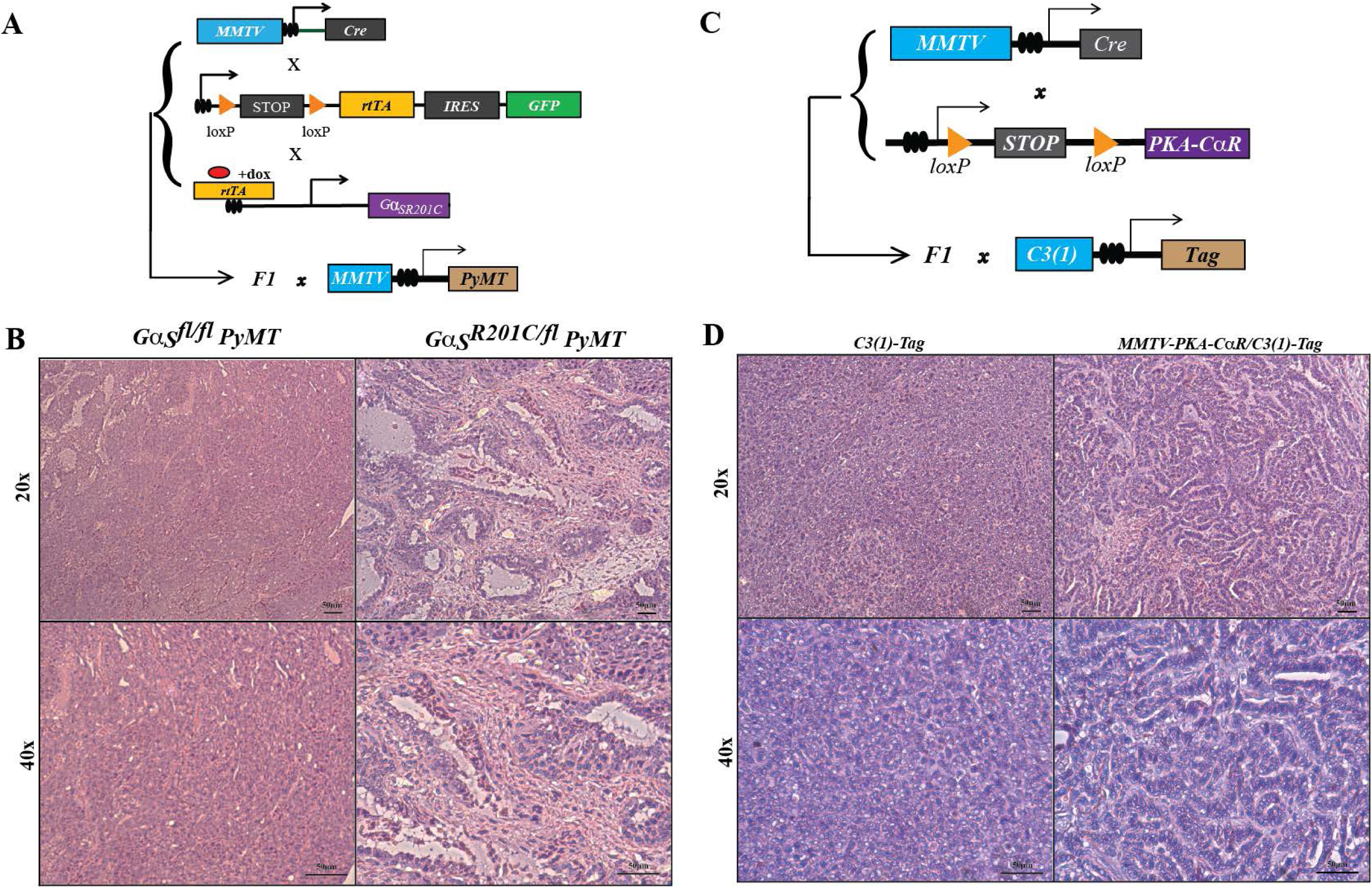
(A) Schematic of mouse crossing strategy to express the doxycycline-inducible *Gα*_*s*_^*R201C/fl*^ allele in the mammary glands of MMTV-PyMT mice following which tumors were harvested and FFPE sections were stained with hematoxylin and eosin (B) to study their histopathology and differentiation status. (C) Schematic of mouse crossing strategy to express the *PKA-CαR* allele in the mammary glands of C3(1)-Tag mice following which tumors were harvested and FFPE sections stained with hematoxylin and eosin to study their histopathology (D).

**Supplementary Figure 4:**
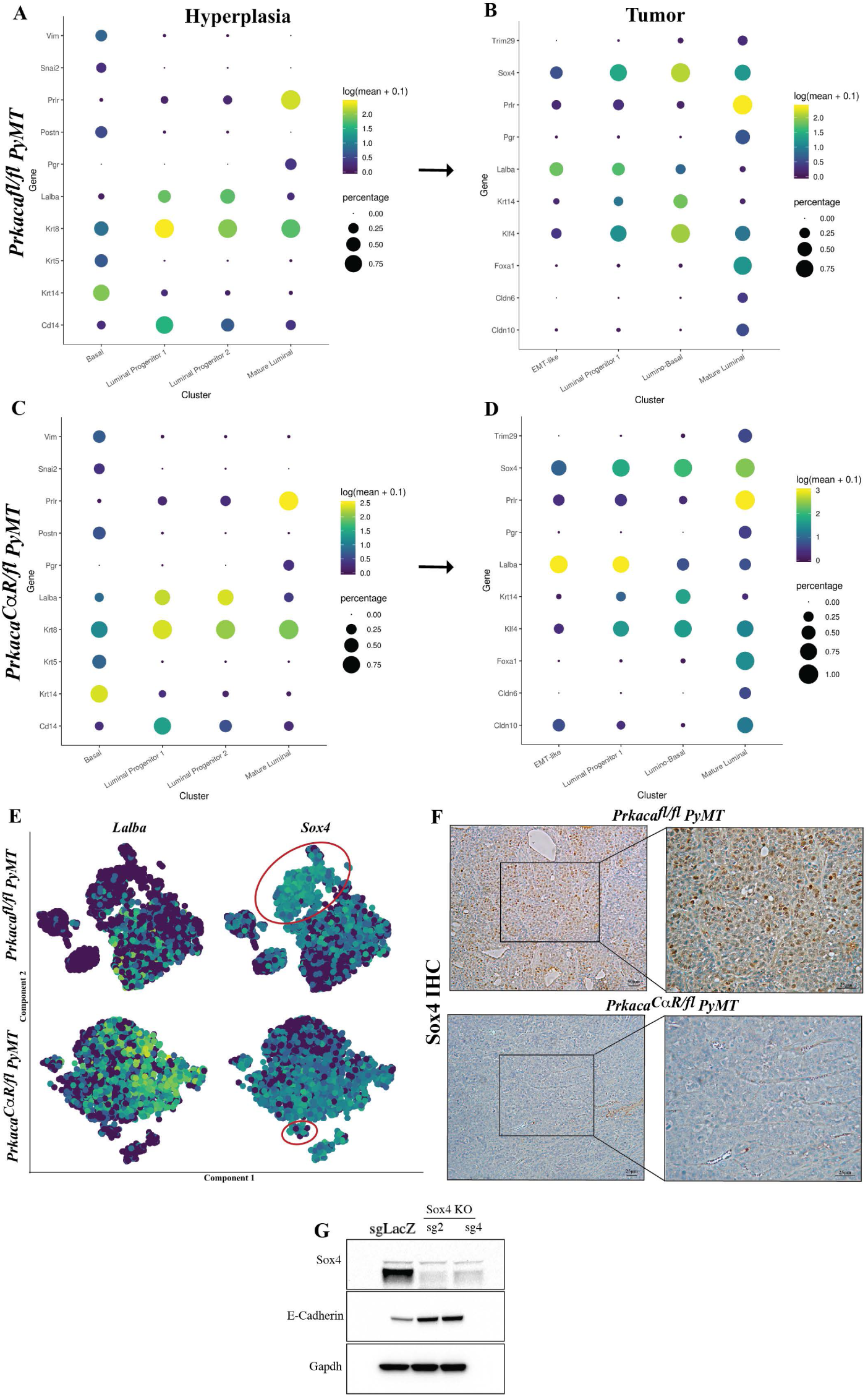
Marker specificity plots outlining the genes whose expression was taken into consideration when annotating each of the subpopulations from 9 week-old hyperplastic glands of *Prkaca*^*CαR/fl*^ PyMT (A) and *Prkaca*^*fl/fl*^ PyMT (C) mice and 22-week old tumors from *Prkaca*^*CαR/fl*^ PyMT (B) and *Prkaca*^*fl/fl*^ PyMT (D) mice. (E) tSNE plots highlighting the expression levels of the *Lalba* (left) and *Sox4* (right) genes in tumors from *Prkaca*^*fl/fl*^ PyMT (top) and *Prkaca*^*CαR/fl*^ PyMT tumors (bottom) The lumino-basal subpopulation is highlighted by the red ellipse. (F) Brightfield images of immunohistochemical staining of Sox4 in FFPE sections from *Prkaca*^*fl/fl*^ PyMT (top) and *Prkaca*^*CαR/fl*^ PyMT (bottom) tumors. (G) Immunoblotting for Sox4, E-cadherin and Gapdh using protein lysates from PB3 cells that contained single guide RNAs targeting LacZ or Sox4.

